# Supply and consumption of glucose 6-phosphate in the chloroplast stroma

**DOI:** 10.1101/442434

**Authors:** Alyssa L. Preiser, Aparajita Banerjee, Nicholas Fisher, Thomas D. Sharkey

**Author notes:** **Author for correspondence:** Thomas D. Sharkey.

## Abstract

Fructose 6-phosphate is an intermediate in the Calvin-Benson cycle and can be acted on by phosphoglucoisomerase to make glucose 6-phosphate (G6P) for starch synthesis. A high concentration of G6P is favorable for starch synthesis but can also stimulate G6P dehydrogenase initiating the glucose-6-phosphate shunt an alternative pathway around the Calvin-Benson cycle. A low concentration of glucose 6-phosphate will limit this futile cycle. In order to understand the biochemical regulation of plastidic glucose 6-phosphate supply and consumption, we characterized biochemical parameters of two key enzymes, phosphoglucoisomerase (PGI) and G6P dehydrogenase (G6PDH). We have found that the plastidic PGI in has a higher *K*_m_ for G6P compared to that for fructose 6-phosphate. The *K_m_* of G6PDH isoform 1 is increased under reducing conditions. The other two isoforms exhibit less redox regulation; isoform 2 is the most inhibited by NADPH. Our results support the conclusion that PGI restricts stromal G6P synthesis limiting futile cycling via G6PDH. It also acts like a one-way valve, allowing carbon to leave the Calvin-Benson cycle but not reenter. We found flexible redox regulation of G6PDH that could regulate the glucose-6-phosphate shunt.

**Highlight:** Glucose 6-phosphate stimulates glucose-6-phosphate dehydrogenase. This enzyme is less active during the day but retains significant activity that is very sensitive to the concentration of glucose 6-phopshate.

## Introduction

Glucose 6-phosphate (G6P) is the first product out of the Calvin-Benson cycle in the starch synthesis pathway. However, it can also enter the oxidative pentose phosphate pathway creating a G6P shunt that bypasses the nonoxidative branch of the pentose phosphate pathway reactions that make up a significant part of the Calvin-Benson cycle. This pathway is generally considered to occur only in the dark (Anderson *et al.*, 1974; Buchanan, 1980; Buchanan *et al.*, 2015; Heldt and Piechulla, 2005; Scheibe *et al.*, 1989). However, a high G6P concentration, favorable for starch synthesis, could cause the shunt to occur in the light. Generally, the G6P concentration in the plastid is low, much lower than its concentration in the cytosol (Gerhardt *et al.*, 1987; Sharkey and Vassey, 1989; Szecowka *et al.*, 2013) but under some conditions the plastid G6P concentration might increase depending on the production and consumption of plastid G6P.

Four different enzymes in the plastid can produce or consume G6P (Fig. 1). First, G6P can be produced by phosphoglucoisomerase (PGI). This enzyme reversibly isomerizes fructose 6-phosphate (F6P) and G6P. Analysis of mutant lines of *Clarkia xantiana* indicated that PGI is not in great excess (Kruckeberg *et al.*, 1989). There are two isoforms of PGI in Arabidopsis, one targeted to the plastid and the other found in the cytosol. The plastid PGI in particular is likely limiting given that G6P/F6P ratios in the plastid are significantly displaced from equilibrium and much lower than in the cytosol (Backhausen *et al.*, 1997; Gerhardt *et al.*, 1987; Schnarrenberger and Oeser, 1974; Sharkey and Vassey, 1989; Szecowka *et al.*, 2013). Plants with loss-of-function mutants in the plastidic enzyme have 98.5% less starch in leaves (Yu *et al.*, 2000). Loss-of-function mutants in the cytosolic enzyme results in increased starch and decreased sucrose (Kunz *et al.*, 2014).

**Fig 1.**
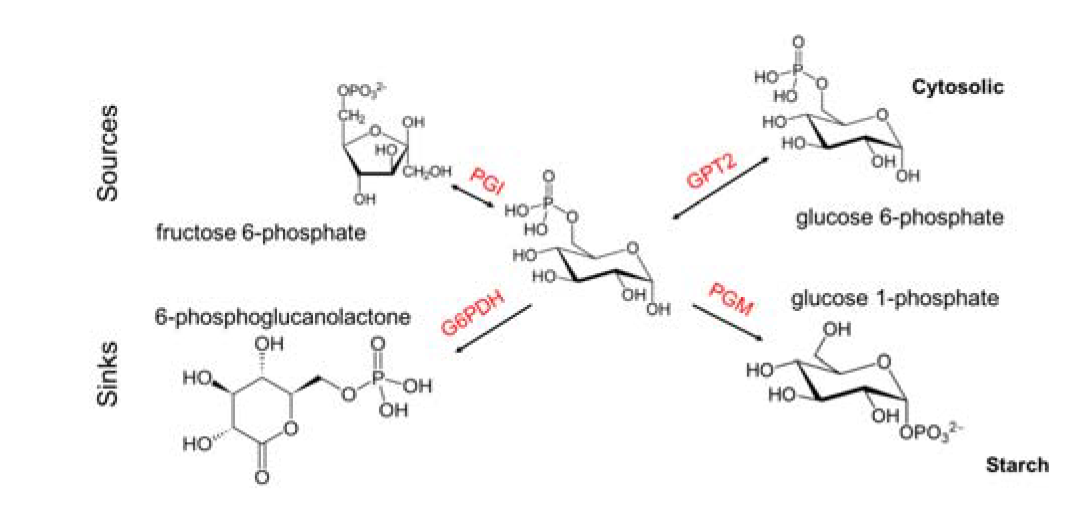
Production and consumption of plastidic G6P. Plastidic G6P can be produced by PGI isomerization of F6P, transported across the plastidic membrane by GPT2, consumed by PGM for starch synthesis, or consumed by G6PDH to enter the G6P shunt.

The second enzyme affecting G6P in the chloroplast is phosphoglucomutase. This enzyme converts G6P to glucose 1-phosphate. This reaction is an important step in starch synthesis. Third, G6P can be transported across the chloroplast membrane by GPT2, a glucose-6-phosphate/phosphate antiporter in the chloroplast membrane. GPT2 is not normally present green tissue (Kammerer *et al.*, 1998; Kunz *et al.*, 2010) and this is corroborated by the large concentration gradient in G6P between the chloroplast and cytosol (Gerhardt *et al.*, 1987; Sharkey and Vassey, 1989; Szecowka *et al.*, 2013). However, GPT2 is important in acclimation to light (Dyson *et al.*, 2015) and is expressed in plants grown in high CO_2_ (Leakey *et al.*, 2009) and is increased when starch synthesis is repressed by knocking out starch synthesis genes (Kunz *et al*. 2010). When GPT2 is present, the gradient of G6P would result in G6P import into the plastid (Gerhardt *et al.*, 1987; Sharkey and Vassey, 1989; Szecowka *et al.*, 2013). Finally, glucose-6-phosphate dehydrogenase (G6PDH) can oxidize G6P to 6-phosphoglucanolactone, the first step in the oxidative branch of the pentose phosphate pathway. There are six isoforms of G6PDH in Arabidopsis. Four of these are predicted to be targeted to the chloroplast where three are functional (Meyer *et al.*, 2011; Wakao and Benning, 2005). It has been hypothesized that during the day, G6PDH initiates a G6P shunt around the Calvin-Benson cycle (Sharkey and Weise, 2016). The G6P shunt oxidizes and decarboxylates G6P to synthesize xylulose 5- phosphate (Xu5P). While the G6P shunt is a futile cycle, it has been proposed to play an important role in stabilization of photosynthesis.

Our goal is to understand the kinetic regulation of the stromal G6P pool, specifically its production by PGI and its consumption by G6PDH. We will not further investigate the roles of PGM and GPT2 since PGM has been characterized because of its key role in starch synthesis (Hattenbach and Heineke, 1999; Najjar, 1948; Ray and Roscelli, 1964), and GPT2 is usually not present in green photosynthetic tissue (Kammerer *et al.*, 1998; Kunz *et al.*, 2010). We repeated critical measurements of PGI kinetics and found that while the isomerization of F6P and G6P is reversible, PGI has a greater affinity for F6P than G6P. Novel findings describing the regulation of G6PDH indicate that G6PDH can remain fairly active during the day. We conclude that a G6P shunt is allowed and even likely in light of the kinetic parameters of G6PDH and that its activity could be modulated during the day to regulate flux through the G6P shunt.

## Materials and Methods

### Overexpression and purification of recombinant enzymes

His-tagged (N-terminal) *Arabidopsis thaliana* plastidic and cytosolic PGI, and C-terminal *Strep*-tagged (Wakao and Benning, 2005; Wendt *et al.*, 2000) plastidic G6PDH1, 2, and 3 genes were commercially synthesized by GenScript (https://www.genscript.com). All of the plasmid constructs were overexpressed in *E. coli* strain BL21. Cells were grown at 37°C to an OD_600_ of 0.6 to1 and induced with 0.5 mM isopropyl β-D-1 thiogalactopyranoside at room temperature. Cells were centrifuged and resuspended in lysis buffer (5 ml lysis buffer/g of pellet; 50 mM sodium phosphate, pH 8.0, 300 mM NaCl) containing 1 mg ml^-1^ lysozyme, 1 μg ml^-1^ of DNAseI, and 1x protease inhibitor cocktail (Sigma, www.sigmaaldrich.com). Cells were then lysed by sonication (Branson Sonifier 250, us.vwr.com). The sonicator was set at 50% duty cycle and an output level of 1. The cells were sonicated using five steps where each step consisted of 15 s pulses and 15 s on ice. The lysate was centrifuged and supernatant collected. For plastidic and cytosolic PGI, Ni-NTA resin (Qiagen, https://www.qiagen.com) was added to the crude lysate with gentle stirring for 1 hr. The mixture was loaded onto a column and washed with wash buffer (50 mM sodium phosphate, pH 8.0, 300 mM NaCl, 10 mM imidazole) until the OD_280_ of the effluent was less than 0.05. Protein was eluted with elution buffer (50 mM sodium phosphate pH 8.0, 300 mM NaCl, 250 mM imidazole) containing 1x protease inhibitor cocktail. The Ni-NTA column purification was performed in a cold room at 4°C. For G6PDH1, 2, and 3, harvested proteins were resuspended in cold Buffer W (IBA, www.iba-lifesciences.com) with protease inhibitor cocktail, 1 mg ml^-1^ lysozyme, and 2.73 kU DNAseI and lysed as described above. Protein was purified on a *Strep-*Tactin column (IBA) following the manufacturer’s instructions. For all purified proteins, SDS-PAGE was carried out and fractions containing >95% of total protein of interest were combined and concentrated using Amicon Ultra 0.5 ml centrifugal filters (molecular weight cut off 3 kDa). Glycerol was added to the concentrated protein to obtain a final protein solution with 15% glycerol. The glycerol stock of the proteins was aliquoted into small volumes, frozen in liquid nitrogen, and stored at −80°C. The concentration of the proteins was determined using Pierce 660 nm protein assay reagent kit (ThermoFisher Scientific, www.thermofisher.com) using a bovine serum albumin standard. Final preparations of purified protein were run on an SDS-polyacrylamide gel and stained with Coomassie Blue to check the purity of the enzymes. Molecular weights were estimated from the protein construct using Vector NTI (ThermoFisher Scientific, www.thermofisher.com).

### Coupled spectrophotometric assay for PGI (F6P to G6P reaction) and G6PDH

The activity of the purified plastidic and cytosolic PGI and G6PDH1, 2, and 3 was studied using coupled spectrophotometric assays. Concentrations of G6P and F6P were determined using NADPH-linked assays measured spectrophotometrically. All assays were validated by demonstrating linear product formation, proportional to the time of the assay and amount of enzyme added. All coupling enzymes were added in excess so that no change in product formation was seen when varying the coupling enzyme. PGI assays were done in 50 mM bicine buffer pH 7.8, containing 4.8 mM DTT, 0.6 mM NADP^+^, 2 U G6PDH (from *Leuconostoc mesenteroides*), varying concentrations of F6P, and 1.31 ng plastidic or cytosolic PGI. The concentrations used to study the *K*_m_ of G6PDH for F6P were 0-4.8 mM. For G6PDH assays, the assay was done in 150 mM Hepes buffer pH 7.2 containing varying concentrations of NADP^+^, varying concentrations of G6P, and G6PDH. 8.3 ng of G6PDH1, 20 ng of G6PDH2, and 44 ng of G6PDH3 were used. The concentrations used to study the *K*_m_ of G6PDH for G6P were 0 – 44.2 mM, and the concentrations used to study *K*_m_ for NADP^+^ were 0 – 11 μM. When G6PDH was assayed varying G6P, 0.6 mM NADP^+^ was added. When G6PDH was assayed varying NADP^+^, 7.6 mM G6P was added for G6PDH1 and 3 and 15.4 mM for G6PDH2. Under these conditions, less than 5% of the non-limiting substrate was consumed over the duration of the assay. The assay mixtures were prepared by adding all the components except the enzyme. Activity was recorded with a dual wavelength filter photometer (Sigma ZFP2) as the change in absorbance at 334 – 405 nm caused by NADP^+^ reduction to NADPH using an extinction coefficient of 6190 M^-1^ cm^-1^. These wavelengths were used because they correspond to mercury lamp emission wavelengths of the lamp used in the filter photometer. When assaying redox sensitivity, G6PDH was incubated with 10 mM DTT or hydrogen peroxide at room temperature before addition to the assay. The G6P concentration was 0.3 mM. For G6P protection assays, G6PDH1 was assayed at 5 mM G6P. After getting a stable baseline with G6PDH and NADP^+^, the reaction was initiated by addition of G6P and incubated at room temperature. Activity was measured 30 or 60 minutes later to allow time for DTT deactivation. Less than 5% of added G6P and NADP^+^ (non-limiting substrate) was consumed within 60 minutes.

### Mass spectrometry assay for PGI (G6P to F6P reaction)

The activity of the purified plastidic and cytosolic PGI in the G6P to F6P direction was studied using a coupled mass spectrometer assay. The assay mixture contained 50 mM Tris pH 7.8, 2.5 mM MgCl_2_, 1 mM ATP, 5 mM DTT, 0.15 U phosphofructokinase (from *Bacillus stearothermophilus*), varying concentrations of G6P, and 1.6 ng of plastidic or cytosolic PGI. The assay mixtures were prepared by adding all the components except the enzyme. The reaction was initiated with the enzyme. After five min, the reaction was quenched with four volumes of 100% ice-cold methanol. Production of FBP was shown to be linear for up to ten min. Five nmol of D-[UL-^13^C_6_] fructose 1,6-bisphosphate was added as an internal standard for quantification, and the sample was heated for 5 minutes at 95°C. Six volumes of 10 mM tributylamine, pH 5.0, was added and the sample was filtered through a Mini-UniPrep 0.2 μm Syringeless Filter Device (GE Healthcare Life Sciences, Whatman). LC/MS-MS was carried out on a Waters Quattro Premier system and was operated in electrospray negative ion mode with both multiple and selected reaction monitoring. The capillary voltage was 2.75 kV; the cone voltage, 50 V; the extractor voltage, 5 V. The source temperature was 120°C and the desolvation temperature was 350°C. Gas flow for the desolvation and cone was set to 800 and 50 l hr^-1^, respectively. The syringe pump flow was 10 μl min^-1^. MassLynx software and the Acquity UPLC Console were used to control the instrument. Samples were passed through an Aquity UPLC BEH Column (Waters) with a multi-step gradient with eluent A (10 mM tributylamine, adjusted to pH 6 with 500 mM acetic acid) and eluent B (methanol): 0-1 min, 95-85% A; 1-3 min, 85%-65% A; 3-3.5 min, 65-40% A; 3.5-4 min, 40-0% A; 4-8.50 min, 0% A; 8.5-10 min, 100% A. The flow rate was 0.3 ml min^-1^. FBP peaks were integrated using MassLynx software and the concentration of the metabolites was quantified by comparing the peak response to a calibration curve.

### Kinetic characterization

Enzymes were assayed at varying concentrations of substrate while keeping the concentration of other substrates (if applicable) constant as described above. The *K*_m_values for plastidic and cytosolic PGI were determined by fitting the data with non-linear regression using the Hill function in OriginPro 8.0 (OriginLab Corporation). All G6PDH isoforms showed substrate inhibition, therefore we estimated regression lines and kinetic constants by finding the minimum of the sum of the squared residuals from the following equation using Solver in Excel, where *v* is the specific activity of the enzyme in μmol mg ^-1^ min^-1^ (Gray *et al.*, 2011):

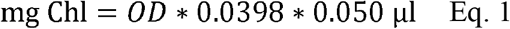

### Inhibition studies

Different metabolites of the Calvin-Benson cycle were tested for their effect on both PGI and G6PDH activity. All the metabolites were purchased from Sigma Aldrich. In metabolite screening assays, metabolites were assayed at a 1:1 ratio with the substrate. To determine the *K_i_* of G6PDH or PGI for different metabolites, the assay was carried out in presence of various concentrations of F6P or G6P and other metabolites. Assay mixtures were prepared as described above with different concentrations of substrate. For PGI studies, 0-0.98 mM F6P or 0-1.5 mM G6P was used and 0-3.8 mM G6P was used for G6PDH studies. The concentration of NADP^+^ in G6PDH assays was held constant at 600 μM. The concentration range used to study the *K_i_* of PGI for E4P were 0-0.05 mM and that for 6PG were 0-1.5 mM. For G6PDH1 and 3 assays 0-0.3 mM NADPH was used and 0-14.5 μM NADPH was used for the G6PDH2 assays. The mechanism of inhibition was determined from Hanes-Woolf plots. The *K_i_* was determined from the non-linear least squares fitting of the activity vs. F6P concentration plot using Solver in Excel using the standard equation for competitive inhibition as described below.

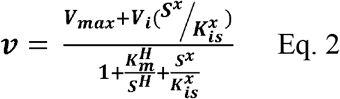

where *V*_max_ is the maximum velocity, *S* is the F6P concentration, *K*_m_ is the Michaelis constant, and *K_i_* is the inhibition constant. For non-competitive inhibition, the below equation was used.

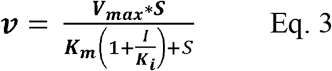

### Midpoint potential of G6PDH1

A series of oxidation-reduction titrations was done with purified G6PDH1. Fully reduced DTT was prepared daily by combining 100 mM DTT with 200 mM sodium borohydride. The mixture was incubated on ice for 20 minutes and then neutralized by adding concentrated HCl to a final concentration of 0.2 M. The mixture was brought to a pH of 8 and diluted to a final concentration of 50 mM DTT. Oxidized DTT and buffers used in the assay were also pH 8. We used mixtures of oxidized and reduced DTT at different redox potentials, ranging from −420 to −124 mV. The total concentration of DTT was 1-8.5 mM. 4.1 ng of G6PDH1 was incubated in the DTT mixture and 1 mg/ml BSA, pH 8 for 1 hr at 25°C in an anaerobic environment. Activity of G6PDH was measured as described in ‘Chloroplast isolation’ with 0.3 mM G6P. The data were fit to the Nernst equation for a two-electron process. We used the *E_m_* of DTT as determined by Hutchison and Ort (1995), −391 mV at pH 8. Oxidized and reduced DTT was quantified using modified protocols from Cho *et al.* (2005) and Charrier and Anastasio (2013) to calculate the potential.

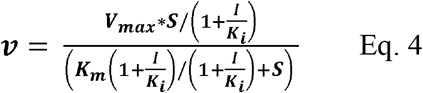

### Leaf extract assays

Wild type *Arabidopsis* was grown in a 12 hr photoperiod at 120 μmol m^-2^ s^-1^ of light. Day temperature was 23°C and night temperature was 21°C. Approximately 300 mg of leaf samples were collected in a 2 ml microfuge tube and immediately frozen by plunging in liquid nitrogen. Samples were ground in a Retsch mill with 4 mm silicone carbide particles (BioSpec Products, www.biospec.com). One ml of extraction buffer (45 mM Hepes, pH 7.2, 30 mM NaCl, 10 mM mannitol, 2 mM EDTA, 0.5% Triton-X-100, 1% polyvinylpolypyrrolidone, 0.5% casein, 1% protease inhibitor cocktail) was added to the sample and vortexed for 30 s. The sample was centrifuged for 30 s at maximum speed and immediately placed on ice. G6PDH activity was assayed as described in “Coupled spectrophotometric assay for phosphoglucoisomerase (F6P to G6P reaction) and G6PDH”. Assays that used leaf extracts were normalized by mg of chlorophyll added to the assay mixture.

### Chloroplast isolation

Fresh spinach was purchased at a local market for use that day. Spinach was either dark or light treated for 1.5 hours before beginning isolation and petioles were kept in water to prevent wilting. Arabidopsis Col-0 was grown on soil in a growth chamber at a 12 h light at 120 μmol m^- 2^s^-1^, 23°C and 12 h dark at 21°C. Plants were harvested either midday for light samples or midnight for dark samples.

Chloroplasts were isolated using a Percoll gradient (Weise *et al.*, 2004). Leaves were placed in a chilled blender with grinding buffer (330 mM mannitol, 50 mM Hepes, pH 7.6, 5 mM MgCl_2_, 1 mM MnCl_2_, 1 mM EDTA, 5 mM ascorbic acid, 0.25% BSA), blended, and then filtered through four layers of cheese cloth. Filtered liquid was centrifuged and the pellet was resuspended in resuspension buffer (330 mM mannitol, 50 mM Hepes, pH 7.6, 5 mM MgCl_2_, 1mM MnCl_2_, 1mM EDTA, 0.25% BSA). The resuspended pellet was layered on top of a 20-80% Percoll gradient which was centrifuged at 1200 g for 7 min. The bottom band in the gradient containing the intact chloroplasts was collected. One volume of resuspension buffer was added to collected chloroplasts and centrifuged at 1200 g for 2 min. The pellet was resuspended in 50 μl of water and vortexed to lyse the chloroplasts. One volume of 2x buffer (100 mM Hepes, pH 7.6, 10 mM MgCl_2_, 2 mM MnCl_2_, 2 mM EDTA, 2 mM EGTA, 60% glycerol, 0.2% Triton X-100, 0.2% PVPP) was added. Samples were stored at −80°C until used for further analysis. Chlorophyll was quantified by lysing 50 μl of purified chloroplasts by sonication and adding supernatant to 1 ml of 95% ethanol. OD_654_ was used to calculate the chlorophyll concentration (Wintermans and DeMots, 1965):

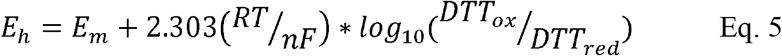

When chloroplast isolations were used to assess activity of fully oxidized and reduced plastidic G6PDH, oxidized or reduced DTT was added at a concentration of 10 mM to each solution used in the isolation. Assays that used isolated chloroplasts were normalized by mg of chlorophyll added to the assay mixture.

## Results

### Purification of recombinant PGI and G6PDH

Final concentration of purified plastidic AtPGI was 15.3 mg/ml and that of cytosolic AtPGI was 13.8 mg/ml (Supplemental Fig. S1). The molecular weight of His-tagged recombinant plastidic and cytosolic AtPGI were ~62.9 kDa and ~62.5 kDa respectively. The specific activity was 787 μmol mg^-1^ protein min^-1^ for plastidic AtPGI and 1522 μmol mg^-1^ protein min^-1^ for cytosolic AtPGI. The final concentration of AtG6PDH1 was 1.66 mg/ml, AtG6PDH2 was 1.90 mg/ml, and AtG6PDH3 was 0.177 mg/ml (Supplemental Fig. S1). The molecular weight of Strep-tagged recombinant AtG6PDH1 was ~65.2 kDa, AtG6PDH2 was ~70.2 kDa, and AtG6PDH3 was 70.5 kDa. The maximum specific activity was 28.0 μmol mg^-1^ protein min^-1^ for AtG6PDH1, 18.7 μmol mg^-1^ protein min^-1^ for AtG6PDH2, and 7.1 μmol mg^-1^ protein min^-1^ for AtG6PDH3.

### Kinetic characterization of plastidic and cytosolic PGI

Table 1 shows the *K*_m_(for both F6P and G6P) of plastidic and cytosolic AtPGI (Supplemental Fig. S2). For plastidic AtPGI, the *K*_m_ value for G6P was ~2.9-fold higher than that for F6P. The *K*_m_’s for F6P and G6P of the cytosolic enzyme were the same. DTT did not significantly influence the specific activity of plastidic or cytosolic PGI (Supplemental Fig. S3).

**Table 1.** Kinetic constants and inhibition constants for plastidic and cytosolic AtPGI as determined by NADPH-linked spectrophotometric assays and LC-MS/MS assays. Each number was determined from the fitted curve as described in the methods.

| | F6P → G6P | | G6P → F6P | |
| --- | --- | --- | --- | --- |
|  | Plastidic PGI | Cytosolic PGI | Plastidic PGI | Cytosolic PGI |
| $K_m$ (μM) | $73 \pm 46$ | $203 \pm 7$ | $164 \pm 25$ | $158 \pm 49$ |
| E4P $K_i$ (μM) | 2.3 | 1.5 | 6.0 | 3.7 |
| 6PG $K_i$ (μM) | 31 | 106 | 245 | 44 |

### E4P and 6PG inhibition of PGI

We tested different metabolites for their effect on PGI activity. Inhibition with either limiting F6P or G6P was similar for both plastidic and cytosolic AtPGI. Erythrose 4-phosphate (E4P), 3-phosphoglyceric acid (PGA), dihydroxacetone phosphate (DHAP), and 6-phosphogluconic acid (6PG) were screened (Fig. 2). Only E4P and 6PG showed significant inhibition. Inhibitory effects were not different between plastidic and cytosolic AtPGI. Fig. 3 shows the activity of plastidic AtPGI over a range of F6P and E4P concentrations. Activity of cytosolic AtPGI was analyzed in a similar manner as shown for plastidic AtPGI (Supplemental Fig. S4). The calculated *K*_i_ values of E4P and 6PG are shown in Table 1. The *K*_i_ values for 6PG were between 31-203 μM, depending on the isoform and substrate. E4P was shown to be more inhibitory with *K*_i_’s between 1.5- 6 μM. Based on the Hanes-Woolf plots (Supplemental Fig. S5), E4P was shown to be competitive, except above 0.04 mM, with G6P. 6PG was identified as competitive with F6P, except above 1.0 mM, and non-competitive with G6P.

**Fig 2.**
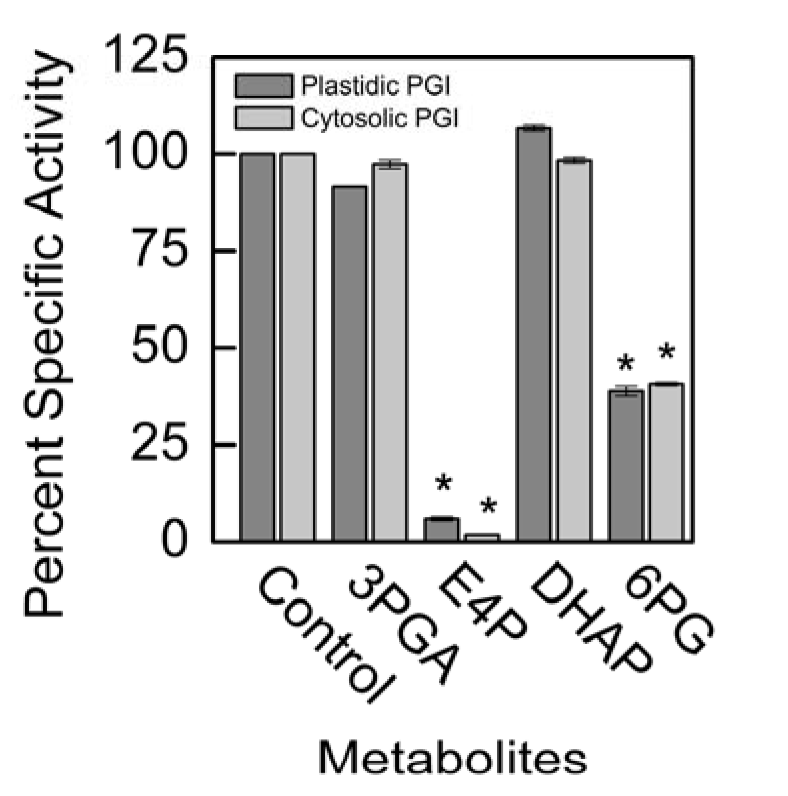
Comparison of specific activity of plastidic (a and b) and cytosolic (c and d) AtPGI with various metabolites. Each bar represents mean and error bars represent S.E. (n=3). All metabolites were screened at 1:1 F6P substrate to metabolite. Data with an asterisk (*) are significantly different from the control as determined by Student’s t-test (P < 0.05).

**Fig 3.**
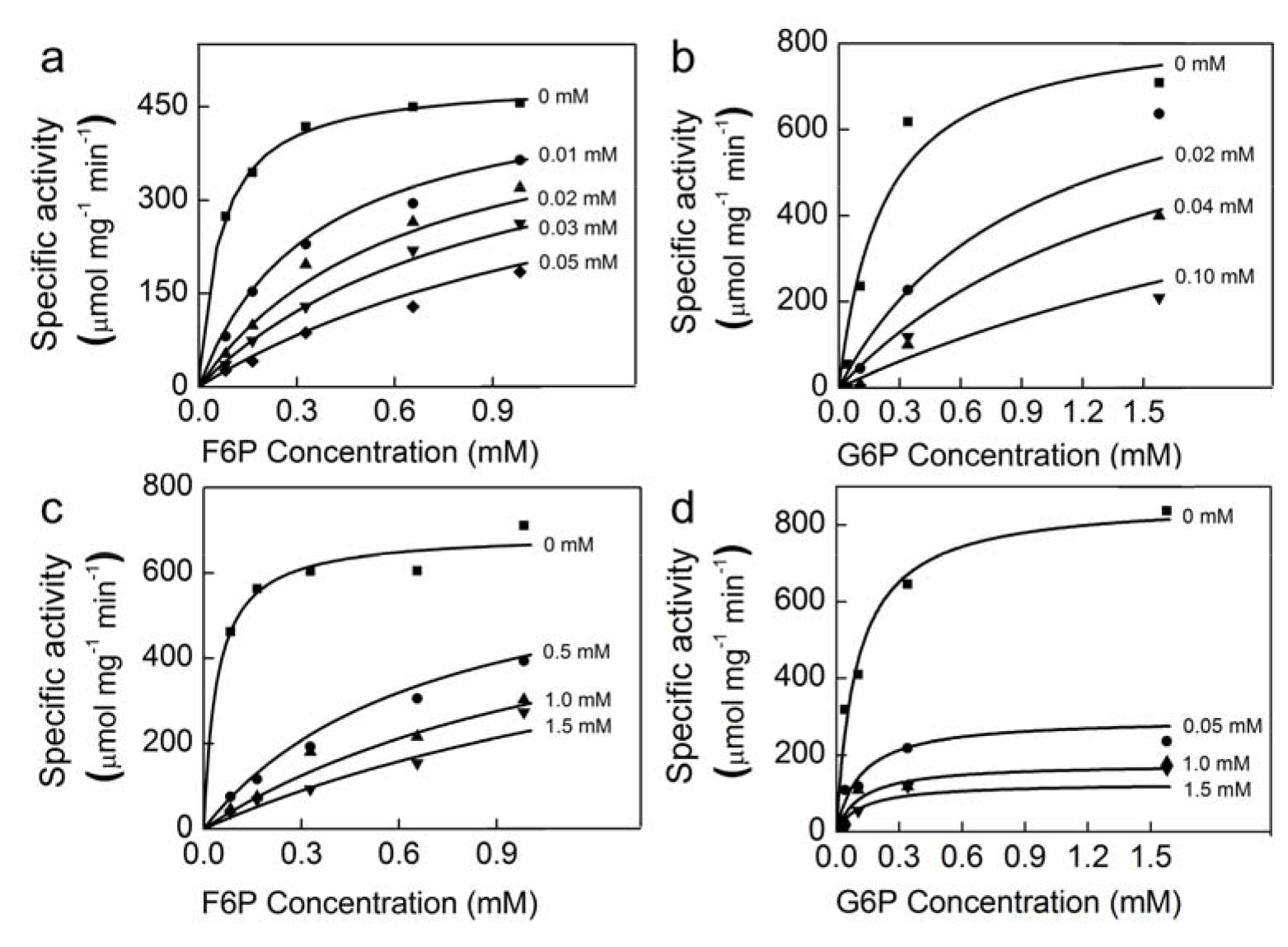
Effect of E4P (a, b) and 6PG (c, d) on plastidic AtPGI. Different symbols represent different concentrations of inhibitor. PGI was more inhibited by E4P than by 6PG. Lines represent data fit to Eq. 3.

### Regulation of PGI in isolated chloroplasts

Plastidic SoPGI activity from chloroplasts from dark-treated spinach leaves had a higher *K*_m_ for G6P compared to light-treated chloroplasts (Fig. 4). The *K_m_* of SoPGI for F6P did not change in the light or dark.

**Fig 4.**
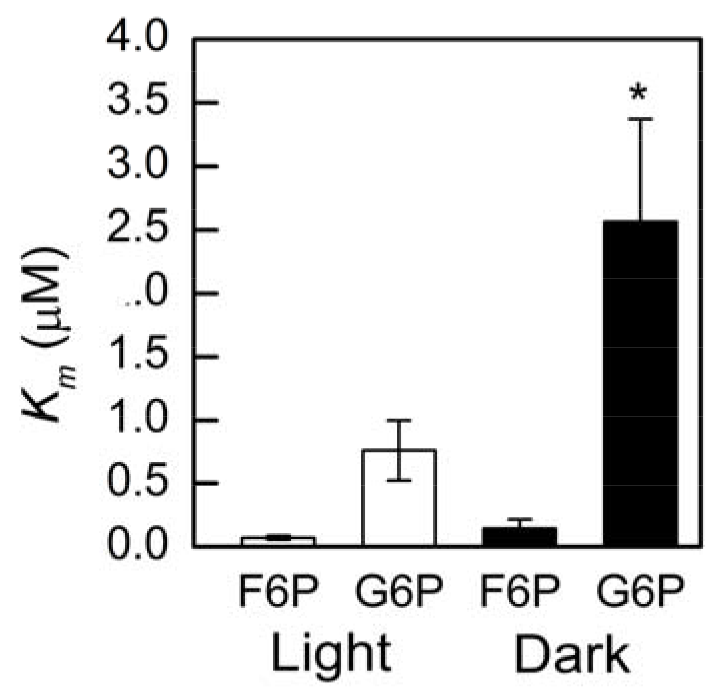
Comparison of G6P and F6P *K_m_* in plastidic SoPGI in dark and light-treated isolated spinach chloroplasts. Each bar represents mean and error bars represent S.E. (n=3). The *K_m_* for G6P increased in dark treated compared to light treated isolated chloroplasts. Bars with a cross (+) are significantly different from corresponding light treated samples as determined by Student’s t-test (P < 0.1).

### Kinetic characterization of G6PDH

All three AtG6PDH isoforms showed substrate inhibition (Fig. 5a). The AtG6PDH *K*_m_ for G6P for isoforms 1 and 3 was 0.3 mM, while the *K*_m_for isoform 2 was approximately 34-fold higher (10.3 mM). Table 2 shows the *K*_m_(for both G6P and NADP^+^), *k*_cat_, and G6P *K*_i_ of all three AtG6PDH isoforms. The catalytic efficiency of AtG6PDH1 for G6P was 190 mM^-1^ s^-1^, of AtG6PDH2 was 3.8 mM^-1^ s^-1^, and of AtG6PDH3 was 48.7 mM^-1^ s^-1^. For NADP^+^, the catalytic efficiency of AtG6PDH1 was 81.4 mM^-1^ s^-1^, of AtG6PDH2 was 30.3 mM^-1^ s^-1^, and of AtG6PDH3 was 29.2 mM^-1^ s^-1^.

**Fig 5.**
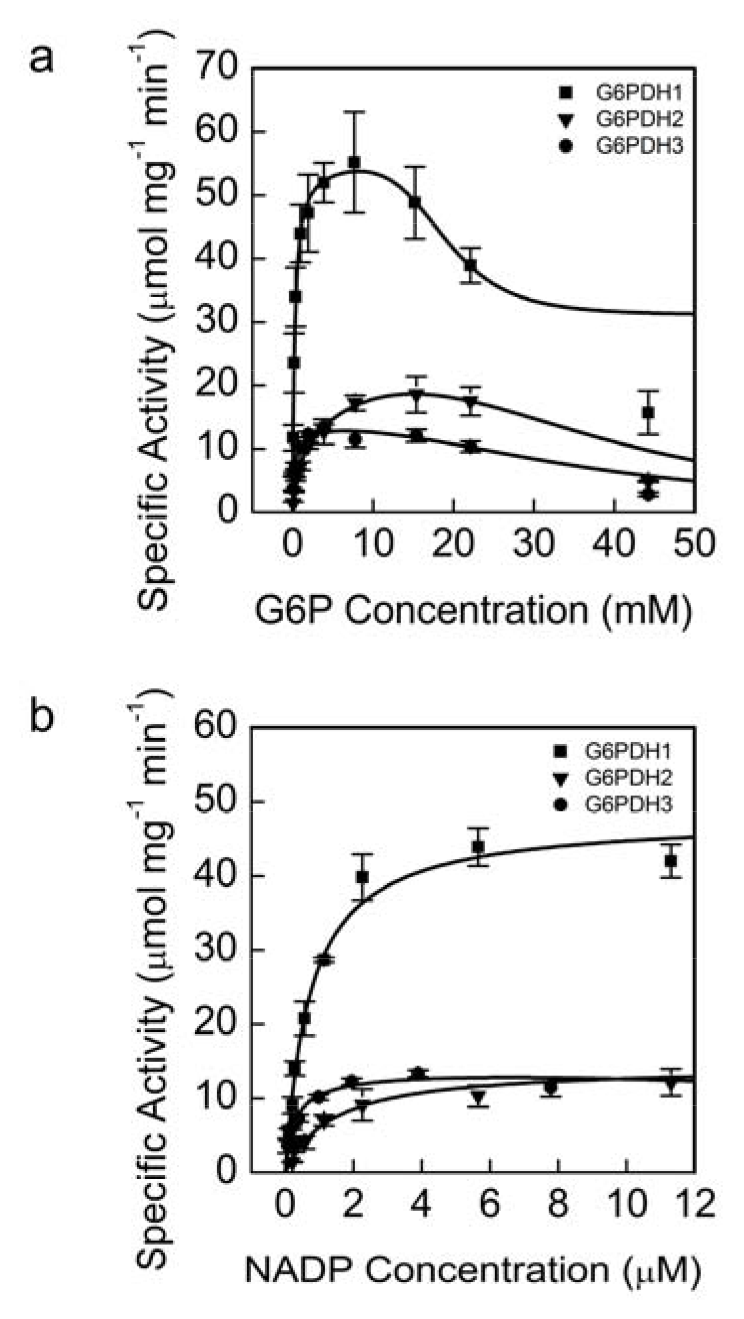
Kinetics of AtG6PDH1, 2, and 3 at different G6P (a) and NADP^+^ (b) concentrations. Each data point represents mean and error bars represent S.E. (n=3). All three isoforms of oxidized G6PDH showed substrate inhibition for G6P. G6PDH1 and 3 showed the greatest affinity for G6P and G6PDH3 had the greatest affinity for NADP^+^. During the G6P experiments NADP^+^ was 0.6 mM and during the NADP^+^ experiment G6P concentration was 7 mM for G6PDH1 and 3 and 15 mM for G6PDH2. In (a) lines represent data fit to Eq. 2 and in (b) lines represent data fit to the Michaelis-Menten equation.

**Table 2.**
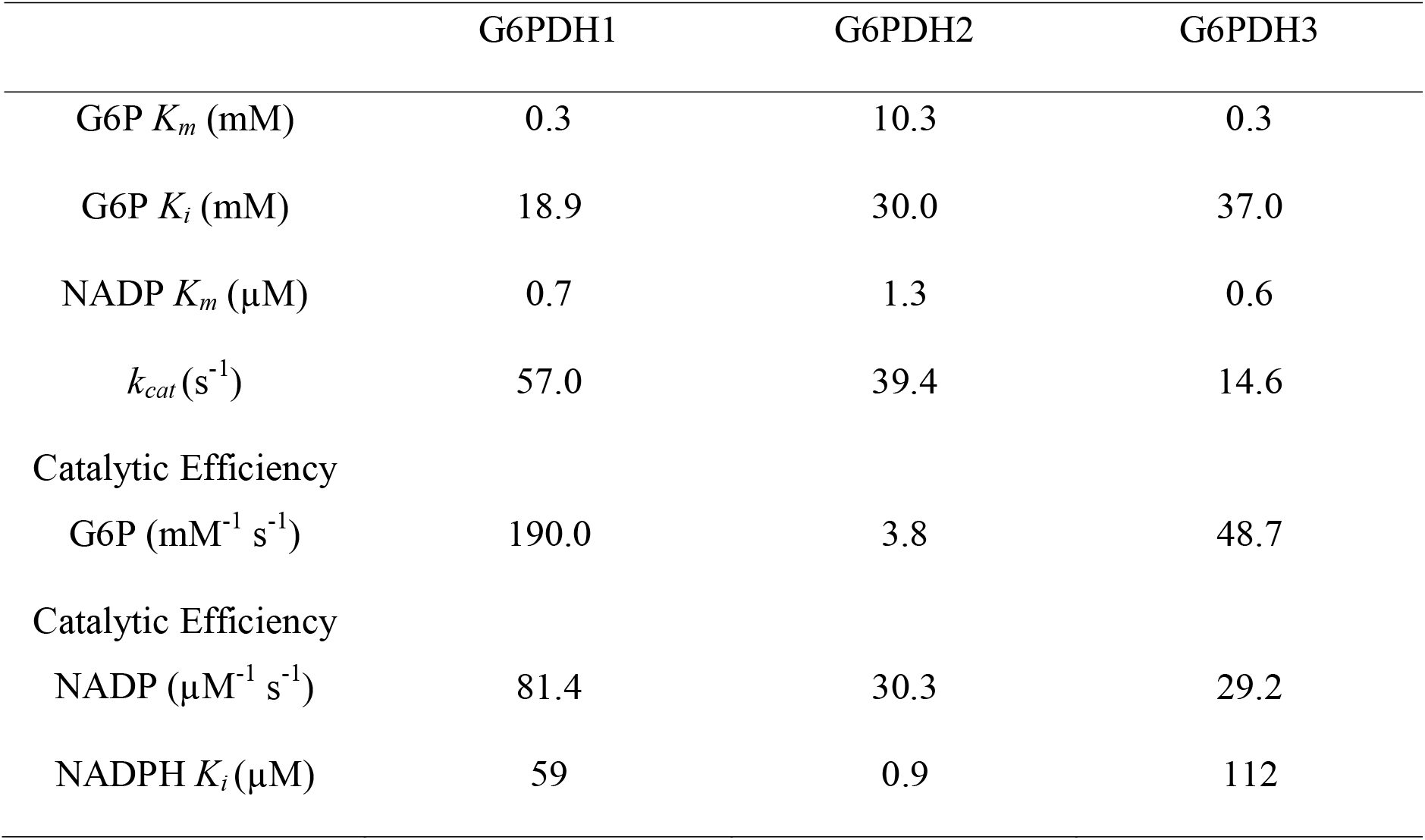
Kinetic constants and inhibition constants of AtG6PDH1, 2, and 3 as determined by NADPH-linked spectrophotometric assays. Each number was determined from by a modified Michalis-Menten equation which includes substrate inhibition. Data points used in model fitting were n=3 different preparations. For inhibition constants, each number was determined from the fitted curves as described in the methods.

### Identification and characterization of inhibitors

We tested ribulose 1,5-bisphosphate (RuBP), ribulose 5-phosphate (Ru5P), F6P, PGA, DHAP, E4P, NADPH, and 6PG for their effect on G6PDH activity. Only NADPH showed inhibition. While NADPH inhibited all three isoforms, AtG6PDH2 was the most inhibited. The calculated *K*_i_ values of NADPH are shown in Table 2. NADPH was found to be competitive for all isoforms based on the Hanes-Woolf plots, except above 14.5 μM for G6PDH2 and above 0.15 mM for G6PDH3 (Supplemental Fig. S5).

### Redox regulation

All isoforms of AtG6PDH were susceptible to deactivation by DTT, but AtG6PDH1 was the most deactivated after two hours, losing approximately 90% of its activity (Fig. 6a). Kinetic characterization of AtG6PDH1 incubated with 10 mM DTT showed that decreased activity in AtG6PDH1 was due to both a decrease in *k*_cat_ and an increase in *K*_m_ and occurred over approximately 45 minutes (Fig. 6b). However, the *k_cat_* was less affected than the *K_m_* (Table 3). Comparison of our results to those of Née *et al.* (2009), who used thioredoxins to deactivate AtG6PDH1, show that DTT is an acceptable mimic of thioredoxins to deactivate AtG6PDH1. Both results show that AtG6PDH1 will lose ~90% of activity when fully reduced. AtG6PDH2 and 3 showed a decrease in *k*_cat_, but not an increase in *K*_m_. AtG6PDH2 retained ~60% of activity and AtG6PDH3 retained ~80% of activity. Redox deactivation of AtG6PDH can be rescued by addition of hydrogen peroxide equimolar to DTT *in vitro* (Fig. 7a). AtG6PDH1 activity reached approximately 64% activity while 79% of the DTT was still reduced (Fig. 7b). The calculated *E_m_* at this time point was −407 mV. Based on our determined midpoint potential of AtG6PDH1 (see Midpoint potential of G6PDH1), we predict AtG6PDH1 would have < 5% activity at the redox potential of the DTT. Therefore, we conclude the addition of hydrogen peroxide did not result in the re-activation of G6PDH1 by oxidizing DTT but that hydrogen peroxide was directly activating G6PDH1. Redox deactivation of G6PDH1 was decreased when G6P was present. When G6P was present at *K_m_* concentration during incubation with DTT, the activity of reduced AtG6PDH was higher than when G6P was not present (Fig. 8).

**Fig 6.**
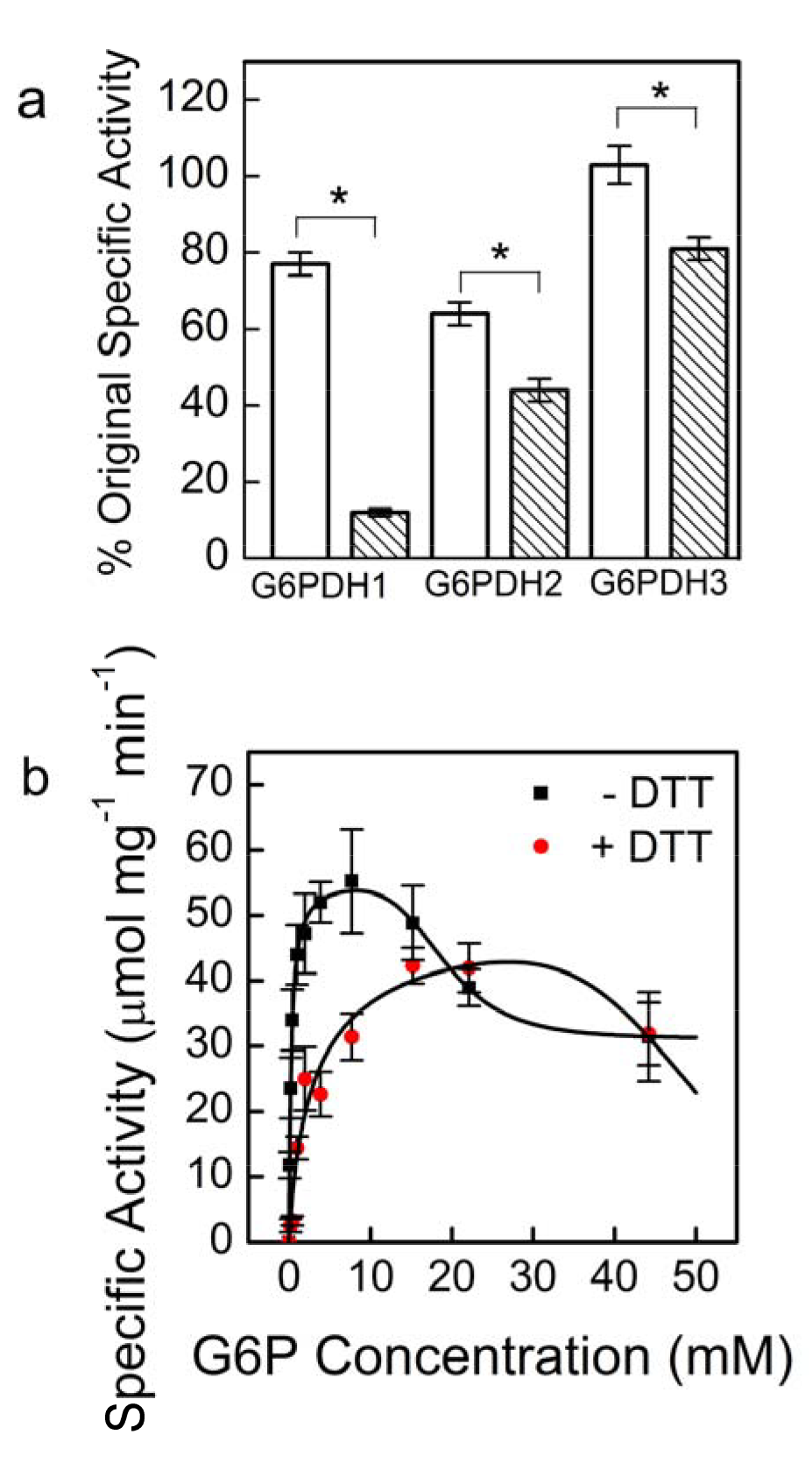
Activity of AtG6PDH 1, 2, and 3 with and without DTT treatment (a) and *K_m_* shift with DTT in G6PDH 1 (b). Each bar or data point represents mean and error bars represent S.E. (n=3). G6PDH1 was the most affected by DTT treatment. White bars indicate controls incubated without DTT for 30 minutes and shaded bars represent incubation with 10 mM DTT for 30 minutes. Assays were done with 5 mM G6P for G6PDH1 and G6PDH3, and 15 mM for G6PDH2. The lines represent data fit to Eq. 2. Bars with an asterisk (*) are significantly different from corresponding controls as determined by Student’s t-test (P < 0.05).

**Fig 7.**
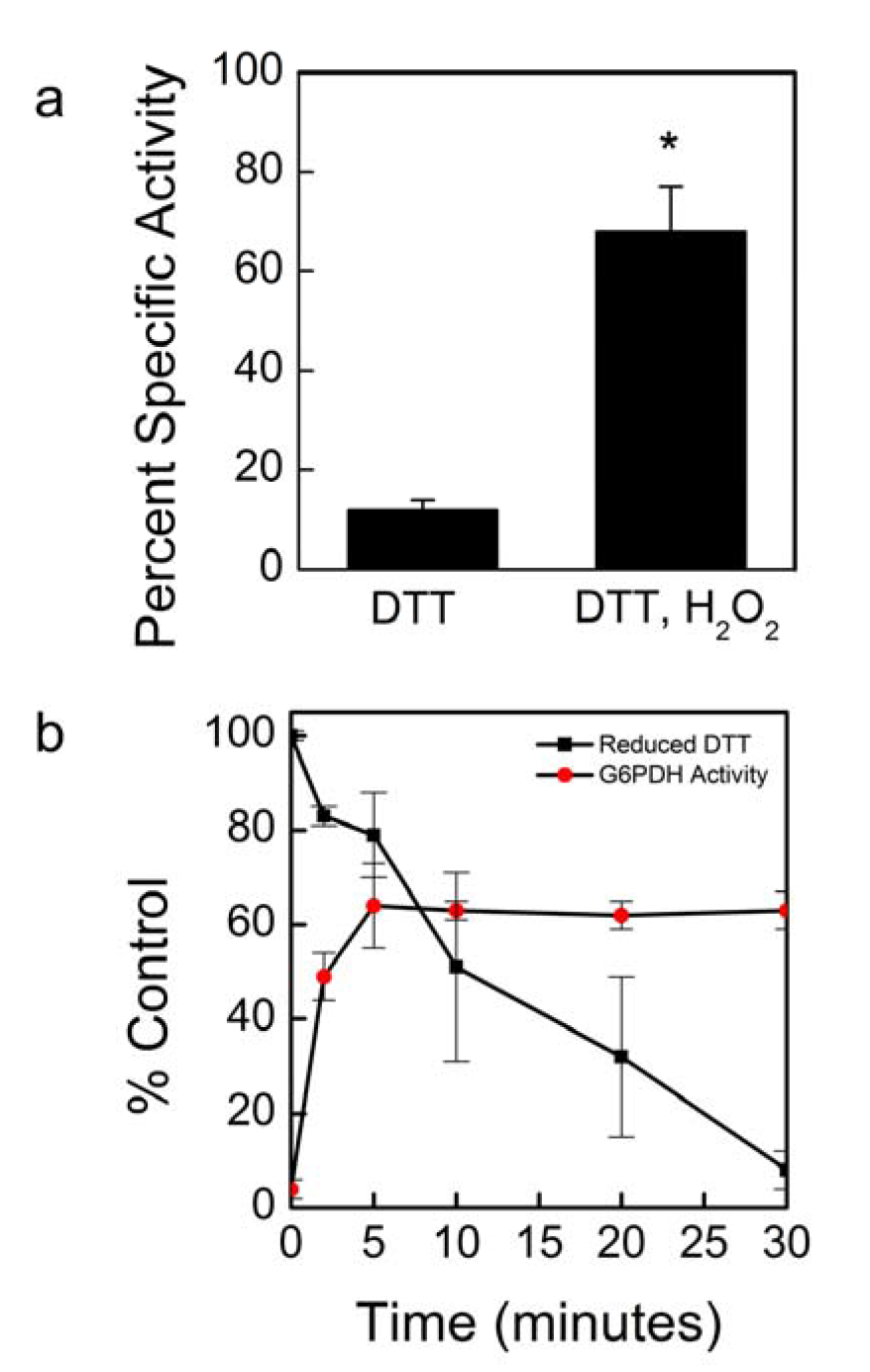
Activation of DTT-deactivated AtG6PDH1 with hydrogen peroxide treatment. G6PDH1 deactivation by DTT could be recovered by addition of equimolar hydrogen peroxide. (a). Reactivation is not through DTT oxidation, but rather hydrogen peroxide directly effects G6PDH1 (b). Assays were done with 5 mM G6P. Each bar or data point represents mean and error bars represent S.E. (n=3). Bars with an asterisk (*) are significantly different as determined by two tailed Student’s t-test (P < 0.05).

**Fig 8.**
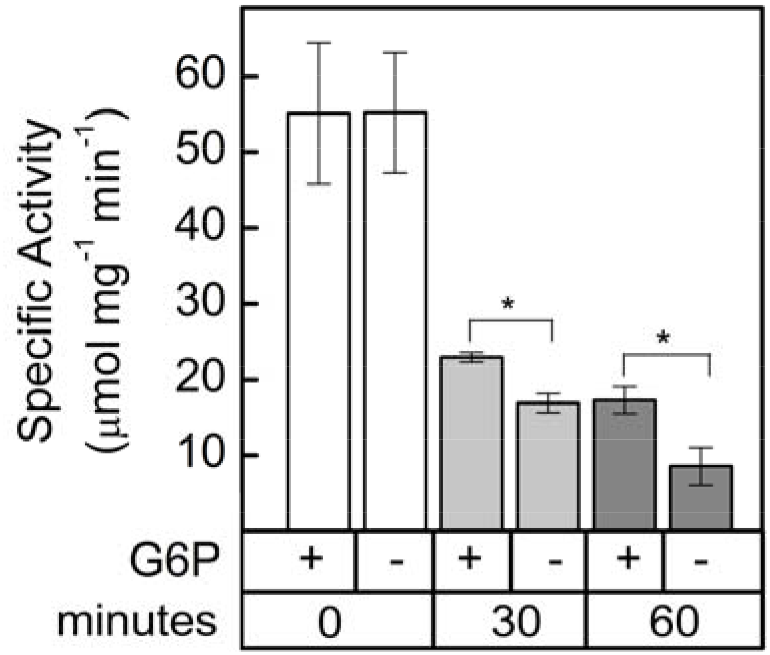
AtG6PDH1 protection from deactivation by G6P. G6PDH1 is less deactivated by DTT after 30 and 60 min when G6P is present at 5 mM. Each bar represents the mean and error bars represent S.E. (n=3). Bars with asterisk (*) are significantly different as determined by two tailed Student’s t-test (P < 0.05).

**Table 3.** Kinetic constants of oxidized and reduced AtG6PDH1, 2, and 3 determined by NADPH-linked spectrophotometric assays. Each number was determined from by a modified Michalis-Menten equation which includes substrate inhibition. Data points used in model fitting were n=3 different preparations.

|  |  | G6PDH1 | G6PDH2 | G6PDH3 |
| --- | --- | --- | --- | --- |
| Oxidized | G6P $K_m$ (mM) | 0.3 | 10.3 | 0.3 |
| | $k_{cat}$ (s<sup>-1</sup>) | 57.0 | 39.4 | 14.6 |
| Reduced | G6P $K_m$ (mM) | 3.4 | 8.6 | 0.6 |
| | $k_{cat}$ (s<sup>-1</sup>) | 52.2 | 20.4 | 11.0 |

### Midpoint potential of G6PDH1

We determined the activity of AtG6PDH1 in a series of oxidation-reduction potentials (Fig. 9). The data was fit with the Nernst equation for a two-electron process. Incubation of AtG6PDH1 at higher redox potentials (−300 to −140 mV) did not increase activity any further. The midpoint potential of G6PDH1 at pH 8 was −378 mV. This corresponds to a midpoint potential of −318 mV at pH 7.

**Fig 9.**
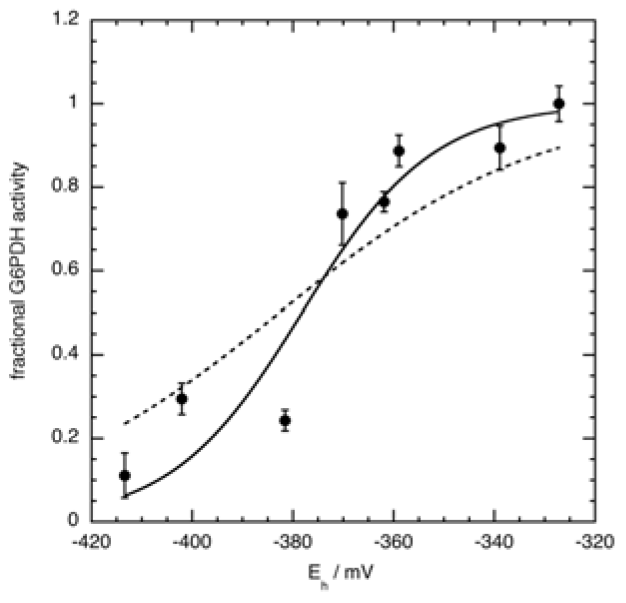
AtG6PDH1 midpoint potential. The midpoint potential of G6PDH1 was determined to be −378 mV at pH 8. Assays were done at the *K_m_* concentration of G6P for oxidized G6PDH1, 0.3 mM. Each data point represents the mean and error bars represent S.E. (n=3). The dashed line represents the Nernst equation for one electron. The solid line represents the Nernst equation for two electrons.

### Activity of G6PDH in isolated chloroplasts and leaf extracts

We used rapid leaf extract assays and chloroplast isolations to determine the activity of redox-regulated SoG6PDH and AtG6PDH compared to total activity in the plastid and the whole leaf. After illumination at 500 μmol m^-2^ s^-1^ for one hr, G6PDH activity spinach leaf extracts decreased by about 35% (Fig. 10a). We also isolated chloroplasts in fully oxidizing or fully reducing conditions. Fully reduced chloroplast activity was about 50% of fully oxidized chloroplast activity in both spinach and *Arabidopsis* (Fig. 10b).

**Fig 10.**
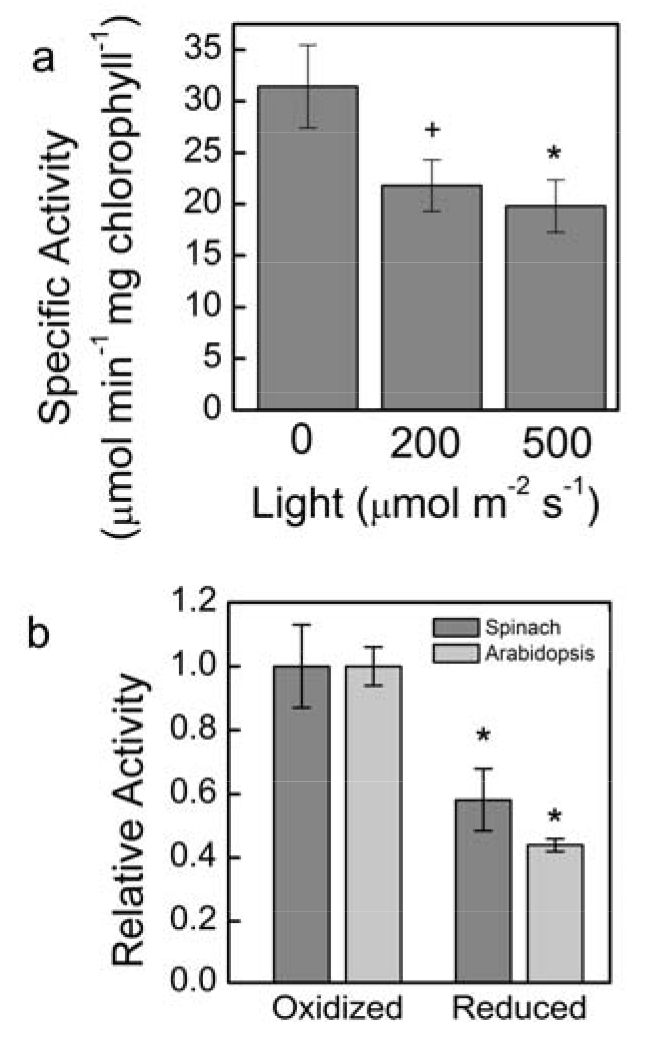
Whole leaf and chloroplast activity of G6PDH in Arabidopsis, and spinach. Whole leaf activity of AtG6PDH decreased 35% after illumination (a). This represents the total redox sensitive G6PDH fraction in Arabidopsis leaves. Chloroplast G6PDH activity decreased 50% after reduction by DTT (b). Samples were normalized by μmol min^-1^ of activity per mg of chlorophyll added to the assay mixture. Assays were done with 5 mM G6P. Each bar represents the mean and error bars represent S.E. (n=3). Bars with a plus sign (+) are significantly different as determined by two tailed Student’s t-test (P < 0.1). One asterisk (*) signifies statistical difference as determined by two tailed Student’s t-test (P < 0.05)

## Discussion

### Regulation of production of G6P

High concentration of G6P in the chloroplast has been proposed to cause a G6P shunt (Fig. 12). The results of this study of key enzymes regulating the stromal G6P concentration support the hypothesis of the G6P shunt.

**Fig 11.**
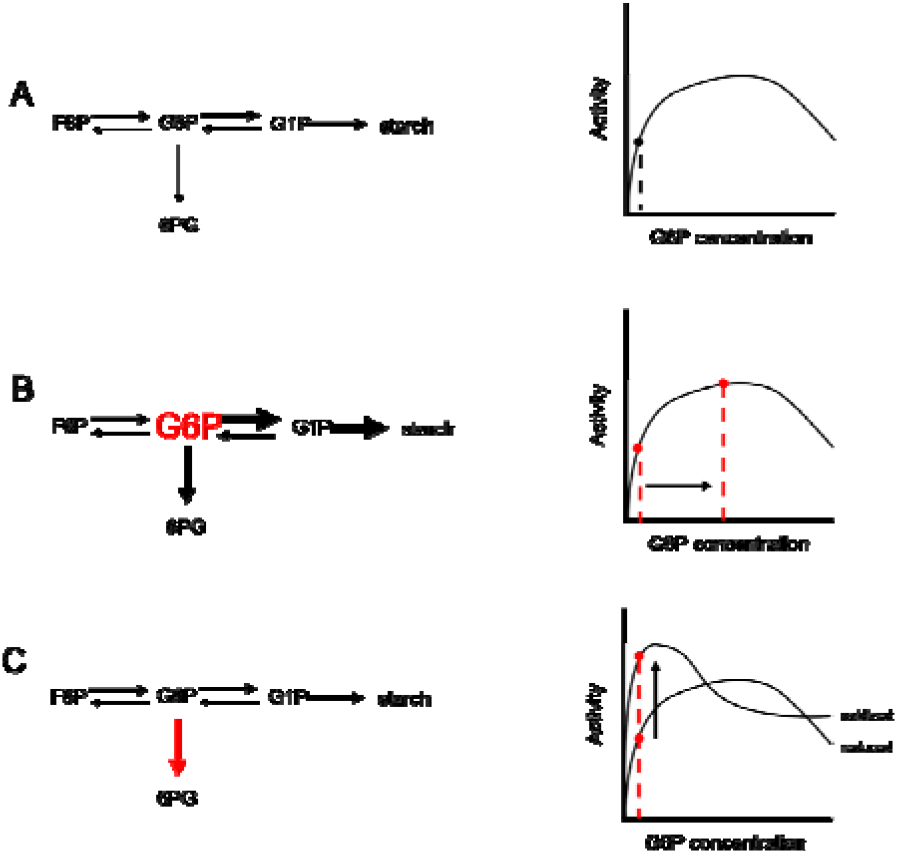
Model of production and consumption of glucose 6-phosphate in the chloroplast stroma through the G6P shunt. Under normal conditions, there may be flux through the G6P shunt as well as flux to starch synthesis (a). Flux through the G6P shunt can by modulated either by an increase in G6P substrate (b) or an increase in G6PDH activity (c). Arrows represent activity of enzymes and changes in thickness represent relative changes in flux. Red represents changes in the steady-state conditions.

**Fig 12.**
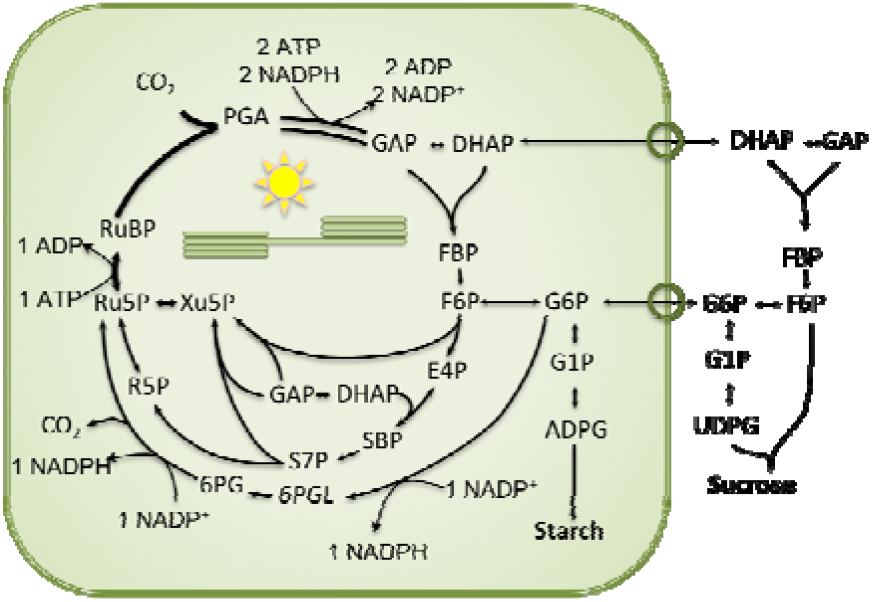
The glucose 6-phosphate shunt. G6PDH is consumed by G6PDH to enter the G6PDH shunt. G6P re-enters the Calvin-Benson cycle as Ru5P. Overall, the shunt consumes three ATP and two NADP^+^ and produces two NAPDH. One CO_2_ molecule is lost for every G6P that enters the shunt.

### PGI

We propose that PGI is a key regulatory point in carbon export from the Calvin-Benson cycle. PGI acts as a one-way valve, going from F6P to G6P. The G6P/F6P ratio at equilibrium has been reported to vary from 3.70 at 10°C to 2.82 at 40°C (Dyson and Noltmann, 1968). However, *in vivo* measurements show the ratio of G6P/F6P in the stroma to be close to 1 (Backhausen *et al.*, 1997; Gerhardt *et al.*, 1987; Schnarrenberger and Oeser, 1974; Sharkey and Vassey, 1989; Szecowka *et al.*, 2013). This disequilibrium is not seen for the cytosolic PGI where G6P/F6P ratios are 2.4-4.7 (Gerhardt *et al.*, 1987; Sharkey and Vassey, 1989; Szecowka *et al.*, 2013). Kinetic hydrogen isotope effects in starch, but not sucrose, also support the conclusion that plastidic PGI, but not cytosolic PGI, has insufficient activity to maintain equilibrium (Schleucher *et al.*, 1999). The high *K*_m_ for G6P for the plastidic enzyme makes this reaction functionally irreversible, helping to explain the kinetic isotope effects only seen in starch. The difference in *K_m_* is seen in both recombinant plastidic Arabidopsis PGI and in isolated plastidic spinach PGI.

We previously assumed that PGA is a strong inhibitor of PGI (eg Sharkey and Weise 2016) based on the report by Dietz (1985). Surprisingly, we did not observe this to be the case. Examination of data from Dietz (1985) shows that during PGA inhibition assays, 6PG was also present in the reaction mixture at 50 μM. The G6P/F6P disequilibrium in chloroplasts was proportional to PGA (Dietz, 1985)(see his Table I) but PGA was not tested alone for its effect on PGI. We found that the *K_i_* of plastidic PGI for 6PG with limiting F6P was 31 μM or with limiting G6P was 203 μM. Based on our findings, we propose that PGI is not inhibited by PGA, and the previously seen inhibition can be explained by presence of 6PG or E4P. *In vivo* plastidic concentrations of 6PG are not known, therefore, extent of inhibition of PGI *in vivo* by 6PG cannot be currently determined.

PGI is inhibited by μM concentrations of E4P (Backhausen *et al.*, 1997; Grazi *et al.*, 1960; Salas *et al.*, 1964). E4P may be inhibitory to both isoforms of PGI because it is a competitive inhibitor and the active sites of both isoforms may be similar (Backhausen *et al.*, 1997). Presumably there is no E4P in the cytosol since it lacks crucial enzymes in the non-oxidative branch of the pentose phosphate pathway (Schnarrenberger *et al.*, 1995). Measurements and estimations of plastidic E4P concentrations *in vivo* show E4P to be ~17-20 μM (Backhausen *et al.*, 1997; Bassham and Krause, 1969; Heldt *et al.*, 1977). This is well above the *K*_i_ of E4P for plastidic PGI. Backhausen *et al.* (1997) propose that this regulation is necessary in order to keep photosynthetic pool sizes stable during changes in light intensity.

In addition to stabilizing the Calvin-Benson cycle, we propose that inhibition of PGI by E4P can provide insight into the phenomenon of reverse sensitivity to CO_2_ and O_2_ of photosynthetic CO_2_ assimilation rate observed by Sharkey and Vassey (1989). They found that when potato leaves were switched to decreased partial pressure of oxygen, rates of photosynthetic CO_2_ assimilation decreased as a result of decreased starch synthesis. Sharkey and Vassey (1989) proposed this was an effect of PGA inhibition of PGI, but because we did not find PGA to be inhibitory, we now suggest that the decrease in starch synthesis is due to an increase in E4P concentrations (or possibly 6PG).

We conclude that PGI is an important regulatory enzyme in central carbon metabolism, keeping G6P concentration lower than would be present at equilibrium thereby regulating the rate of the G6P shunt but also starch synthesis. Overexpression of phosphoglucomutase significantly increased starch synthesis confirming that starch synthesis is regulated at PGI in addition to the well-known regulation at ADPglucose pyrophosphorylase (Uematsu *et al.*, 2012).

### GPT2

If GPT2 is present it can import G6P from the cytosol to the chloroplast. Niewiadomski *et al.* (2005) showed that GPT2 could restore starch accumulation to plants lacking PGI. Expression of GPT2 is present in leaves when starch synthesis is blocked by loss of starch synthesis enzymes, when plants are grown in high CO_2_, or exposed to an increase in light intensity (Dyson *et al.*, 2015; Kunz *et al.*, 2010; Leakey *et al.*, 2009). It also is expressed in CAM plants, which require high rates of starch synthesis (Cushman *et al.*, 2008; Neuhaus and Schulte, 1996). Thus, we believe that much of the time plants rely on PGI alone to supply G6P for starch synthesis and regulate PGI to regulate the supply of G6P to control the rate of the shunt. When higher rates of starch synthesis are needed GPT2 is expressed, increasing the supply of G6P (Fig. 11) but making the plant vulnerable to high rates of the G6P shunt.

### Regulation of consumption of G6P by G6PDH

Stromal G6P is primarily thought of as an intermediate in starch synthesis. It is converted by phosphoglucomutase to glucose 1-phosphate. However, there are additional reactions involving stromal G6P can participate in in the plastid. Here, we investigated consumption of G6P by G6PDH. G6PDH is competitively inhibited by its product NADPH and redox regulation that results mostly in an increase in *K*_m_, which reduces futile cycling in leaves in the light (Scheibe *et al.*, 1989; Wakao and Benning, 2005). However, while the enzyme is less active in the light (Anderson *et al.*, 1974; Buchanan, 1980; Buchanan *et al.*, 2015; Heldt and Piechulla, 2005; Scheibe *et al.*, 1989) our results show that it retains significant activity. Two factors that will modulate G6PDH activity in the light are the sensitivity of G6PDH to G6P and the redox regulation of G6P.

### Sensitivity to G6P

The activity of G6PDH (and thus the G6P shunt) is sensitive to stromal G6P concentration by three mechanisms.

- Deactivation of G6PDH in reducing conditions is primarily due to an increase in *K_m_* (Scheibe *et al.*, 1989, Fig 6).
- G6P protects G6PDH from deactivation in reducing conditions (Fig 8).
- G6P has been shown to relieve the inhibition of G6PDH by NADPH, as well as decrease the *K_m_* and increases the *k_cat_* of G6PDH in assays where NADP is varied (Olavarría *et al.*, 2012; Shreve and Levy, 1980).

In conditions where G6P concentrations in the stroma may increase, such as those discussed above in “Production of G6P”, flux through G6PDH (and the G6P shunt) would also increase.

### Redox Regulation

We have determined the midpoint potential of G6PDH to be −378 mV at pH 8. This agrees with results from Née *et al.* (2009) and is close to the midpoint potential of other redox regulated enzymes in the Calvin-Benson cycle and electron transport (Cammack *et al.*, 1977; Hirasawa *et al.*, 1998; Hirasawa *et al.*, 2000; Hirasawa *et al.*, 1999; Knaff, 2000; Née *et al.*, 2009; Strand *et al.*, 2016). Assuming equilibrium and the midpoint potential of G6PDH1 at pH 8 as a reference, using the Nernst equation, we calculate that all Calvin-Benson enzymes and electron transport proteins are almost fully reduced and thus active while G6PDH maintains 50% of its activity (Table 4). Exceptions are ferredoxin and malate dehydrogenase (MDH), which are predicted to be oxidized at −378 mV. Although there may be deviations from redox equilibrium within the stroma, from these approximations we conclude that the midpoint potential of AtG6PDH1 is in a range to allow dynamic regulation of G6PDH and that it is theoretically possible to have flux through the Calvin-Benson cycle and the G6P shunt at the same time.

**Table 4.**
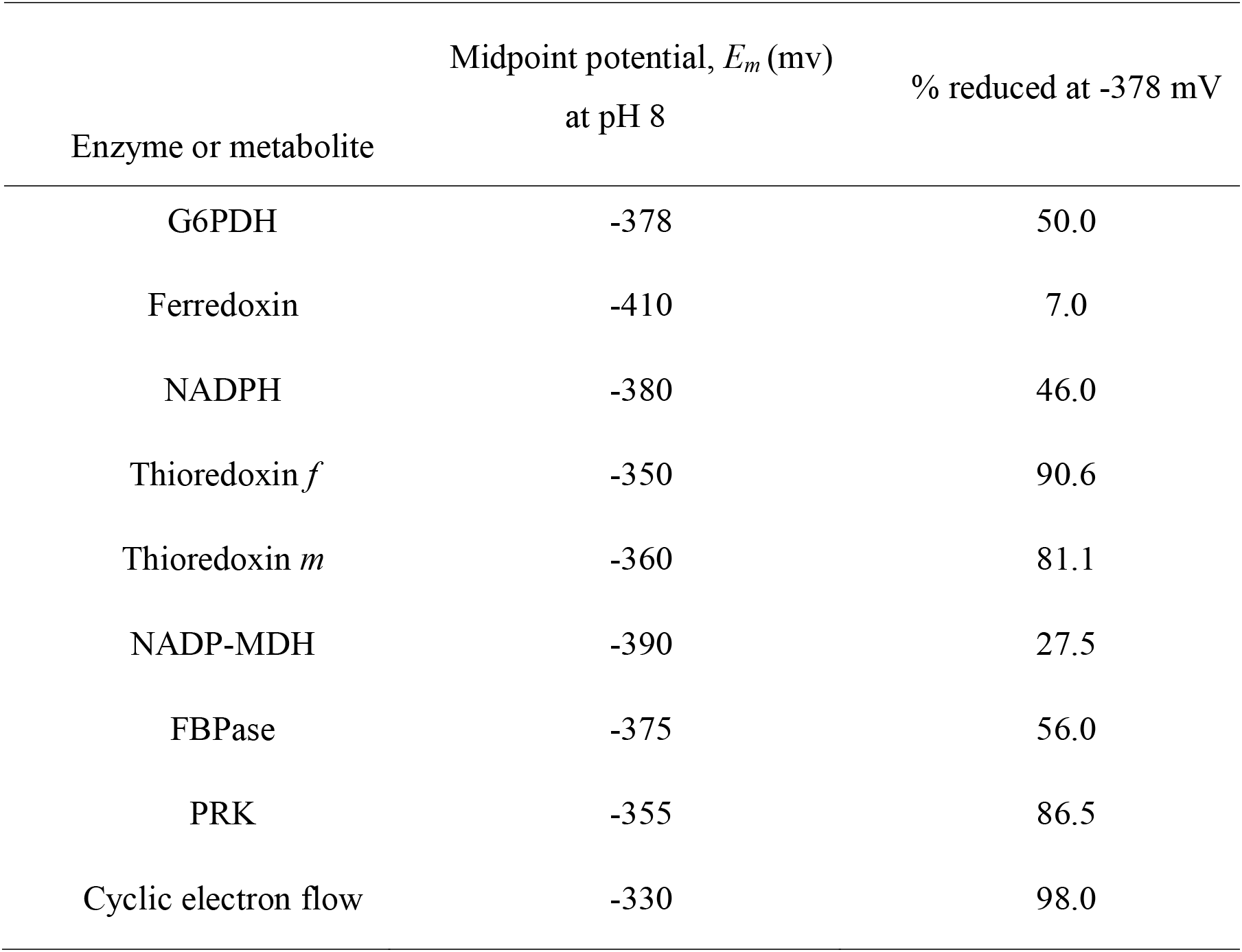
Midpoint potentials and percent reduction of key Calvin-Benson cycle enzymes and electron transport proteins at −378 mV at pH 8, assuming equilibrium. Calvin-Benson cycle enzymes are mostly active at the midpoint potential of G6PDH. One electron chemistry is assumed for ferredoxin and two electron chemistry for all others.

| Enzyme or metabolite | Midpoint potential, $E_m$ (mv)<br>at pH 8 | % reduced at -378 mV |
| --- | --- | --- |
| G6PDH | -378 | 50.0 |
| Ferredoxin | -410 | 7.0 |
| NADPH | -380 | 46.0 |
| Thioredoxin *f* | -350 | 90.6 |
| Thioredoxin *m* | -360 | 81.1 |
| NADP-MDH | -390 | 27.5 |
| FBPase | -375 | 56.0 |
| PRK | -355 | 86.5 |
| Cyclic electron flow | -330 | 98.0 |

We have also shown that G6PDH can be activated upon addition of hydrogen peroxide. Brennan and Anderson (1980) and Née *et al.* (2009) previously demonstrated a role for hydrogen peroxide regulation of G6PDH both *in vivo* and *in vitro* in the presence of thioredoxin. In conditions where hydrogen peroxide can accumulate, such as high light, G6PDH deactivation could be reversed to modulate the consumption of G6P by the G6P shunt. The activity of G6PDH can be modulated by redox status of the plastid, G6P concentration, and hydrogen peroxide.

Redox regulation of dominant isoforms of G6PDH is found in many species, including Arabidopsis, pea, potato, spinach, and barley (Scheibe *et al.*, 1989; Schnarrenberger *et al.*, 1973; Semenikhina *et al.*, 1999; Wenderoth *et al.*, 1997; Wendt *et al.*, 2000; Wright *et al.*, 1997). We have shown that plastidic G6PDH from isolated Arabidopsis chloroplasts retains approximately 50% of its total activity, even in high light conditions. Additionally, cytosolic G6PDH is not redox regulated and makes up 33% of whole leaf G6PDH activity. G6P in the cytosol could be converted to pentose phosphate and be imported into the plastid by the XPT transporter (Eicks *et al.*, 2002).

Oxidative stress might also stimulate the G6P shunt. Drought or high light can result in an accumulation of hydrogen peroxide and other ROS products (see Suzuki *et al.* (2011) for a review). Based on current findings, we propose that, with accumulation of hydrogen peroxide, the *K*_m_ of G6PDH1 can decrease, increasing the flux through the G6P shunt. Sharkey & Weise (2016), proposed that the shunt can induce cyclic electron flow, which may help protect PSI. Photoprotective mechanisms of PSII, for example state transitions of the antenna complex or energy dependent quenching, are usually sufficient to safely dissipate excess excitation energy at PSII (Derks *et al.*, 2015). However, with high light, in fluctuating light (Allahverdiyeva *et al.*, 2014), and at low temperature (Sonoike, 2011), excess energy or electrons could still be passed on to PSI and result in PSI photoinhibition. Unlike PSII, the proteins of PSI have a low turnover rate and damage to PSI is considered more severe (Scheller and Haldrup, 2005; Sonoike, 2011). Coupling ATP consumption in the G6P shunt with cyclic electron flow would dissipate light energy at PSI (Miyake *et al.*, 2004; Munekage *et al.*, 2004; Strand and Kramer, 2014).

### Conclusion

Our data supports the conclusion that production and consumption of plastidic G6P is carefully regulated. Plastidic PGI activity is not adequate to bring F6P and G6P to equilibrium, preventing an accumulation of G6P, and G6PDH is partially deactivated reducing loss of carbon while still maintaining regulatory flexibility to increase and decrease the G6P shunt (Fig. 12) as needed.

## Supplementary data

**Supplemental Fig. S1**- SDS page of the purified G6PDH and PGI proteins, stained with Coomassie blue.

**Supplemental Fig. S2**- Kinetics of plastidic and cytosolic AtPGI at different F6P and G6P concentrations.

**Supplemental Fig. S3**- Effect of 10 mM DTT on plastidic and cytosolic AtPGI

**Supplemental Fig. S4**- Hanes-Woolf plots of E4P and 6PG inhibition of plastidic AtPGI

**Supplemental Fig. S5-** Hanes-Woolf plots of NADPH effect on G6PDH1

## Acknowledgements

We thank Michigan State University Research Technology Support Facility Mass Spectrometry Core for providing the facility for doing the LC-MS/MS work. This research was funded by U.S. Department of Energy Grant DE-FG02-91ER2002 (T.D.S. and A.L.P) and DE-FG02- 11ER16220 (N.F.). A.L.P is partially supported by a fellowship from Michigan State University under the Training Program in Plant Biotechnology for Health and Sustainability (T32- GM110523). Partial salary support for T.D.S. came from Michigan AgBioResearch.

## Author Contributions

A.L.P. designed and carried out the experiments and analyzed the data. A.B. designed and carried out the PGI experiments. N.F. did the calculations for the midpoint potential experiments. A.L.P. wrote the manuscript. T.D.S. supervised the project and edited the manuscript. All authors discussed the results and provided critical feedback.

## Abbreviations

6PG: 6-phosphogluconic acid
At: *Arabidopsis thaliana*
DHAP: dihydroxacetone phosphate
E4P: erythrose 4-phosphate
F6P: fructose 6-phosphate
FBP: fructose 1,6-bisphosphate
G6P: glucose 6-phosphate
G6PDH: glucose-6-phosphate dehydrogenase
PGA: 3-phosphoglyceric acid
PGI: phosphoglucoisomerase
PGM: phosphoglucomutase
So: Spinacia oleracea
Xu5P: xylulose 5-phosphate

